# GREB1 amplifies androgen receptor output in prostate cancer and contributes to antiandrogen resistance

**DOI:** 10.1101/433755

**Authors:** Eugine Lee, John Wongvipat, Danielle Choi, Ping Wang, Deyou Zheng, Philip A. Watson, Anuradha Gopalan, Charles L. Sawyers

## Abstract

Genomic amplification of the androgen receptor (AR) is an established mechanism of antiandrogen resistance in prostate cancer. Here we show that the magnitude of AR signaling output, independent of AR genomic alteration or expression level, also contributes to antiandrogen resistance, through upregulation of the coactivator GREB1. We demonstrate 100-fold heterogeneity in AR output within cell lines and show that cells with high AR output have reduced sensitivity to enzalutamide. Through transcriptomic and shRNA knockdown studies, together with analysis of clinical datasets, we identify GREB1 as a gene responsible for high AR output. We show that GREB1 is an AR target gene that amplifies AR output by enhancing AR DNA binding and promoting p300 recruitment. GREB1 knockdown in high AR output cells restores enzalutamide sensitivity in vivo. Thus, GREB1 is a candidate driver of enzalutamide resistance through a novel feed forward mechanism.

## Introduction

Androgen receptor (AR) targeted therapy is highly effective in advanced prostate cancer but is complicated by the emergence of drug resistance, called castration-resistant prostate cancer (CRPC) (Shen & Abate-Shen, 2010; Watson, Arora, & Sawyers, 2015). The most common mechanism of CRPC is restored AR signaling, primarily through amplification of AR (C. D. Chen et al., 2004; Robinson et al., 2015). The importance of AR amplification as a clinically important drug resistance mechanism is underscored by recent data showing that AR amplification, detected in circulating tumor DNA or in circulating tumor cells (CTCs), is correlated with reduced clinical benefit from the next generation AR inhibitors abiraterone or enzalutamide (Annala et al., 2018; Podolak et al., 2017).

Genomic landscape studies of prostate cancer have revealed several molecular subtypes defined by distinct genomic drivers (Berger et al., 2011; Cancer Genome Atlas Research, 2015; Taylor et al., 2010). In addition to this genomic heterogeneity, primary prostate cancers also display heterogeneity in AR transcriptional output, measured by an AR activity score (Hieronymus et al., 2006). Notably, these differences in transcriptional output occur in the absence of genomic alterations in AR, which are generally found only in CRPC (Cancer Genome Atlas Research, 2015). One potential explanation for this heterogeneity in AR transcriptional output is through coactivators and other AR regulatory proteins such as FOXA1, SPOP, FOXP1 and TRIM24 (Cancer Genome Atlas Research, 2015; Geng et al., 2013; Groner et al., 2016; Pomerantz et al., 2015; Takayama et al., 2014).

Much of the work to date has focused on inter-tumoral heterogeneity. Here we address the topic of intra-tumoral heterogeneity in AR transcriptional output, for which we find substantial evidence in prostate cancer cell lines and in primary prostate tumors. Using a sensitive reporter of AR transcriptional activity to isolate cells with low versus high AR output, we show that high AR output cells have an enhanced response to low doses of androgen and reduced sensitivity to enzalutamide, in the absence of changes in AR mRNA and protein expression. To understand the molecular basis for these differences, we performed transcriptome and shRNA knockdown studies and identified three genes (GREB1, KLF8 and GHRHR) upregulated in high AR output cells, all of which promote AR transcriptional activity through a feed-forward mechanism. Of these, we prioritized GREB1 for further characterization because GREB1 mRNA levels are increased in primary prostate tumors that have high AR activity. GREB1 amplifies AR transcriptional activity through a two-part mechanism: by promoting p300 recruitment and by enhancing AR binding to chromatin. Importantly, GREB1 knockdown converted high AR output cells to a low AR output state and restored enzalutamide sensitivity in vivo. Collectively, these data implicate GREB1 as an AR signal amplifier that contributes to prostate cancer disease progression and antiandrogen resistance.

## Results

### Isolation of cells with low and high AR output but comparable AR expression

Previous work using a PSA promoter/GFP reporter (PSAP-eGFP) showed that LNCaP prostate cancer cells display varying levels of eGFP expression. Characterization of low GFP cells in this analysis revealed reduced AR levels and increased expression of stem cell and developmental gene sets (Qin et al., 2012). We explored this question in the context of the contemporary data on heterogeneity in AR transcriptional output using a different AR-responsive reporter, ARR_3_tk-eGFP, where eGFP expression is driven by the probasin promoter modified to contain three AR responsive elements (Snoek et al., 1998). LNCaP (Figure 1) and CWR22PC-EP (Figure 1-figure supplement 1) prostate cancer cells containing a single copy of the reporter construct were derived by infection with lentivirus containing the reporter at a low multiplicity of infection (MOI) (Figure 1A). Remarkably, we observed >100-fold range in eGFP expression, as measured by flow cytometry, despite similar levels of AR by immunofluorescence microscopy (Figure 1B,C, Figure 1-figure supplement 1A).

**Figure 1.**
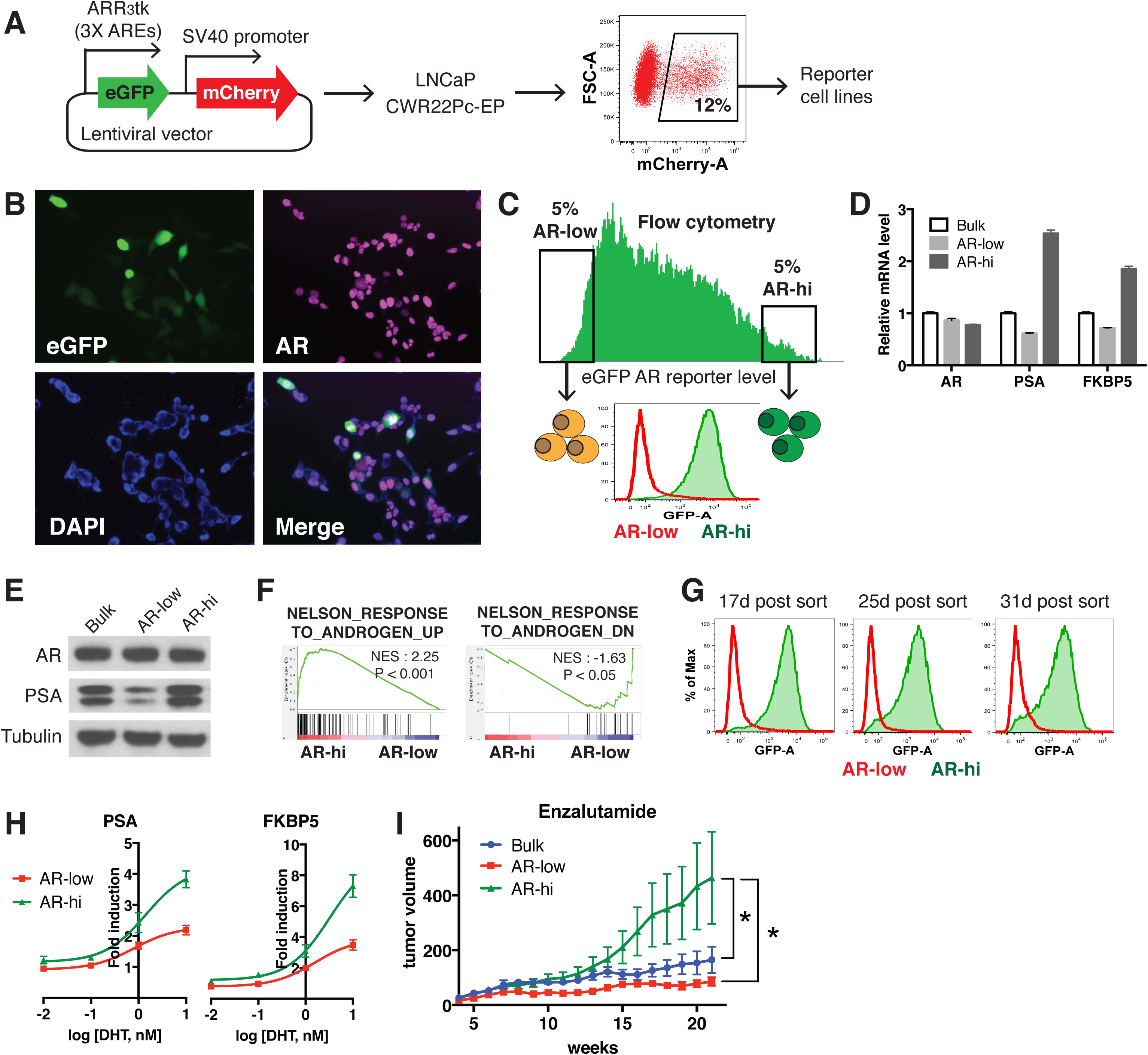
Characterization of LNCaP prostate cancer cells with low vs. high AR output. (**A**) The reporter cell lines were generated by infecting the cells with the lentivirus containing eGFP AR reporter construct (details can be found in methods). Cells stably integrated with the construct were sorted out by mCherry flow cytometry. (**B**) The variable AR reporter activity (green) in LNCaP cells. Cells were also stained with AR (magenta) and DAPI (blue). (**C**) LNCaP cells with low (AR-low) and high (AR-hi) AR activities were sorted out using flow cytometry based on eGFP AR-reporter expression. (**D-E**) AR-hi cells have higher AR output while having same level of AR. The q-PCR data (D) presented as mean fold change ± SD relative to bulk population. (**F**) Gene set enrichment analysis (GSEA) shows that the gene sets up- and down-regulated by androgen are enriched in AR-hi and AR-low cells, respectively. (**G**) AR-low and AR-hi cells maintain their AR activities over time. (**H**) AR-hi cells showed enhanced up-regulation of AR target genes in response to DHT treatment. The q-PCR data presented as mean fold change ± SD relative to DMSO control. (**I**) LNCaP/AR xenografts derived from AR-hi cells become resistant to enzalutamide faster than other populations. The bulk, sorted AR-low and AR-hi cells were injected into castrated mice and the mice were treated with enzalutamide immediately after injection. Data presented as mean ± SEM (N=10). *P<0.05 (One-way ANOVA).

We then used flow cytometry to isolate eGFP-positive cells from both ends of the spectrum of AR transcriptional output, which we refer to as AR-hi (high AR output) and AR-low (low AR output) cells respectively (Figure 1C, Figure 1-figure supplement 1A). AR-hi cells also express higher levels of endogenous AR target genes (FKBP5, PSA, TRPM8) (Figure 1D,E, Figure 1-figure supplement 1B,C), and have an overall increase in AR transcriptional activity based on RNA-sequencing analysis (Figure 1F). In addition, the AR-low and AR-hi transcriptional phenotypes remain stable for over 30 days post sorting (Figure 1G, Figure 1-figure supplement 1D). Interestingly, AR-low cells showed upregulation of gene sets related to proliferation and cell cycle (Figure 1-source data 1). Of note, Qin et al (Qin et al., 2012) reported downregulation of these gene sets in their low/absent PSA cells, suggesting that the two reporters read out different transcriptional activities. Importantly, the difference in AR output between AR-low and AR-hi cells is not explained by different levels of AR expression or nuclear translocation, since both were comparable in each subpopulation (Figure 1D,E, Figure 1-figure supplement 1B,C, Figure 1-figure supplement 2).

We next asked if isolated AR-low and AR-hi populations have different responses to ligands such as dihydrotestosterone (DHT) or antagonists such as enzalutamide. AR-hi cells showed enhanced sensitivity to DHT in a dose-dependent manner (Figure 1H; Figure 1-figure supplement 1E). This result is similar to the effect of increased AR expression in conferring sensitivity to low doses of androgen (C. D. Chen et al., 2004), but now without a change in AR level. To address sensitivity to enzalutamide, we used LNCaP/AR xenografts (derived from LNCaP cells) because this model has a track record of revealing clinically relevant mechanisms of enzalutamide resistance (Arora et al., 2013; Balbas et al., 2013). As we did with LNCaP and CWR22PC-EP cells, we derived AR-low and AR-hi subpopulations by flow cytometry and also observed differential AR output despite similar levels of AR expression (Figure 1-figure supplement 3A-C). Remarkably, AR-hi cells developed enzalutamide resistance significantly faster that AR-low or parental cells when injected into castrated mice treated with enzalutamide (Figure 1I).

Having demonstrated heterogeneous AR output within prostate cancer cell lines, we asked if similar, intra-tumoral heterogeneity is observed clinically by immunohistochemical analysis of PSA and AR expression in several primary cancers. Consistent with previous reports (Qin et al., 2012; Ruizeveld de Winter et al., 1994), we observed heterogeneous PSA staining that is not strictly correlated with AR level. For example, we found variable intensity of PSA staining in tumor cells with comparable levels of AR staining (lined boxes; Figure 1-figure supplement 4) and, conversely, variable intensity of AR staining in tumor cells with similar PSA staining (dotted circles; Figure 1-figure supplement 4). Although this is a small dataset, the results indicate that the AR transcriptional heterogeneity we observe in prostate cancer cell lines is present in patient samples. Emerging technologies for conducting single cell RNA and protein analysis in clinical material will enable deeper investigation of this question.

### GREB1 maintains high AR transcriptional output

To elucidate the molecular basis underlying the differences in AR-low and AR-hi cells, we performed RNA-sequencing and found 69 genes upregulated in AR- low cells and 191 genes upregulated in AR-hi cells (fold change ≥ 1.5, p < 0.05, Figure 2-source data 1). In addition to enrichment of gene sets regulated by androgen (Figure 1F), human prostate luminal and basal cell gene sets were enriched in AR-hi and AR-low cells, respectively (Figure 2A). Based on these results we postulated that high AR output could be a consequence of upregulation of transcriptional co-activators and/or of genes involved in luminal differentiation. We therefore filtered the list of 191 genes upregulated in AR-hi cells and identified 33 genes annotated as co-activators or luminal genes (Figure 2-source data 2), then measured the consequence of shRNA knockdown of each one on AR output in AR-hi cells (Figure 2B). 3 of the 33 candidate genes (GREB1, GHRHR, KLF8) inhibited AR activity when knocked down in AR-hi cells, with successful knockdown confirmed by qRT-PCR (Figure 2C,D). AR knockdown served as a positive control, and ACPP (one of the 30 genes that did not score) served as a negative control. Interestingly, all three hits are transcriptional upregulated by DHT simulation (Figure 2E), which likely explains their increased expression in AR-hi cells.

**Figure 2.**
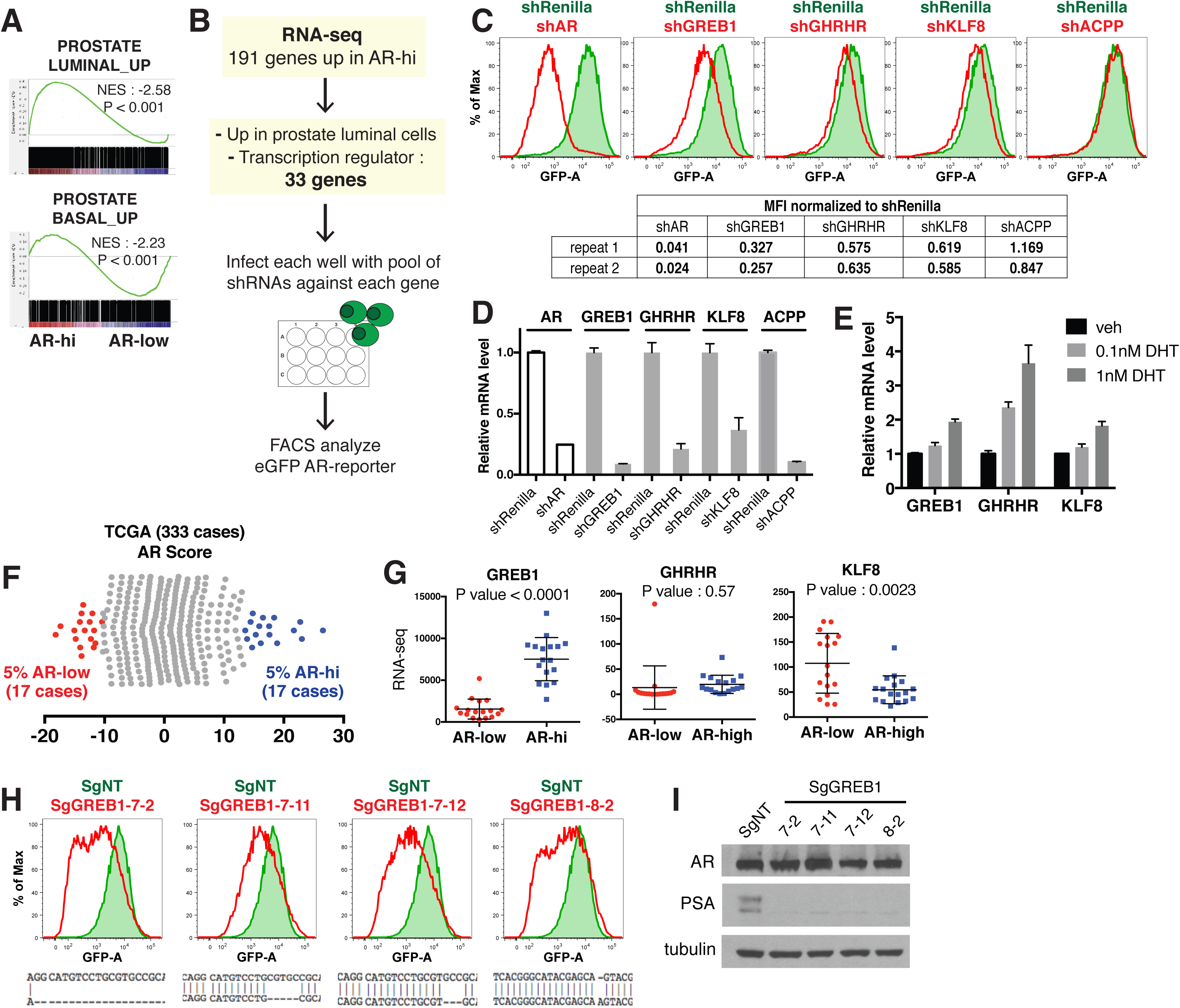
Knockdown of the three AR regulated genes, GREB1, GHRHR and KLF8, inhibited AR activity in cells with high AR activity. (**A**) Gene set enrichment analysis (GSEA) shows that genes upregulated in human prostate luminal and basal cells are enriched in AR-hi and AR-low cells, respectively. (**B**) The schematic of knockdown study with 33 selected genes upregulated in AR-hi cells. Details can be found in methods. (**C**) The flow cytometry results show that the knockdown of GREB1, GHRHR and KLF8 inhibited AR reporter activity. Top: The flow cytometry plot of one of the duplicate assays is shown. Bottom: The normalized median fluorescence intensity (MFI) of eGFP reporter in each assay is shown. AR shRNA was used as a control. ACPP knockdown result was shown as a representative hairpin that had no affect on reporter activity. (**D**) The knockdown level of AR, GREB1, GHRHR, KLF8 and ACPP in (C). The q-PCR data presented as mean fold change ± SD relative to shRenilla control. (**E**) The transcription of GREB1, GHRHR and KLF8 is regulated by androgen. The data presented as mean fold change ± SD relative to DMSO control. (**F**) Graph showing AR score of each TCGA primary prostate tumor. Red and blue points are tumors with lowest (AR-low) and highest (AR-hi) AR score, respectively (5% of 333 cases: 17 cases each). (**G**) The RNA levels of GREB1, GHRHR and KLF8 are compared between AR-low vs. AR-hi TCGA cases (data represent mean ± SD, unpaired t-test). (**H**) The GREB1 function is inhibited by CRISPR/Cas9 in four LNCaP sublines. (Top) AR reporter activity is inhibited in all four GREB1 CRISPR cell lines compared to control (SgNT). (Bottom) The example of genomic alteration in target sequence at each cell line is shown. (**I**) The CRISPR/Cas9-mediated inhibition of GREB1 suppressed PSA expression without affecting AR level.

Among the three, GREB1 emerged as the most compelling candidate for further investigation based on interrogation of clinical datasets. Specifically, we found increased expression of GREB1, but not GHRHR or KLF8, in primary prostate tumors from the TCGA dataset with high AR output scores (top 5%) versus low AR output scores (bottom 5%) (Figure 2F,G). To be sure that GREB1 is relevant in other model systems, we confirmed GREB1 upregulation in CWR22PC-EP AR-hi cells (Figure 2-figure supplement 1A) and reduced AR output after GREB1 knockdown (Figure 2-figure supplement 1B). We further validated the knockdown data using CRISPR/Cas9, which also showed inhibition of AR output (by flow cytometry) and highly reduced PSA expression in LNCaP AR-hi sublines expressing different sgRNAs targeting GREB1, without detectable changes in AR protein level (Figure 2H,I).

### GREB1 amplifies AR transcriptional activity by enhancing AR DNA binding

GREB1 was first reported as an estrogen-regulated gene in breast cancer (Rae et al., 2005) then shown to bind directly to ER, presumably through its LxxLL motif, and function as an ER coactivator by promoting interaction with cofactors (Mohammed et al., 2013). To determine if GREB1 also functions as an AR coactivator, we introduced exogenous GREB1 (HA-GREB1) into AR-low LNCaP and CWR22PC-EP cells and derived stably expressing sublines (Figure 3A, Figure 3-figure supplement 1A). GREB1 overexpression enhanced DHT-induced AR target gene expression in a dose-dependent manner (Figure 3B,C, Figure 3-figure supplement 1B), indicating that GREB1 also promotes AR activity.

**Figure 3.**
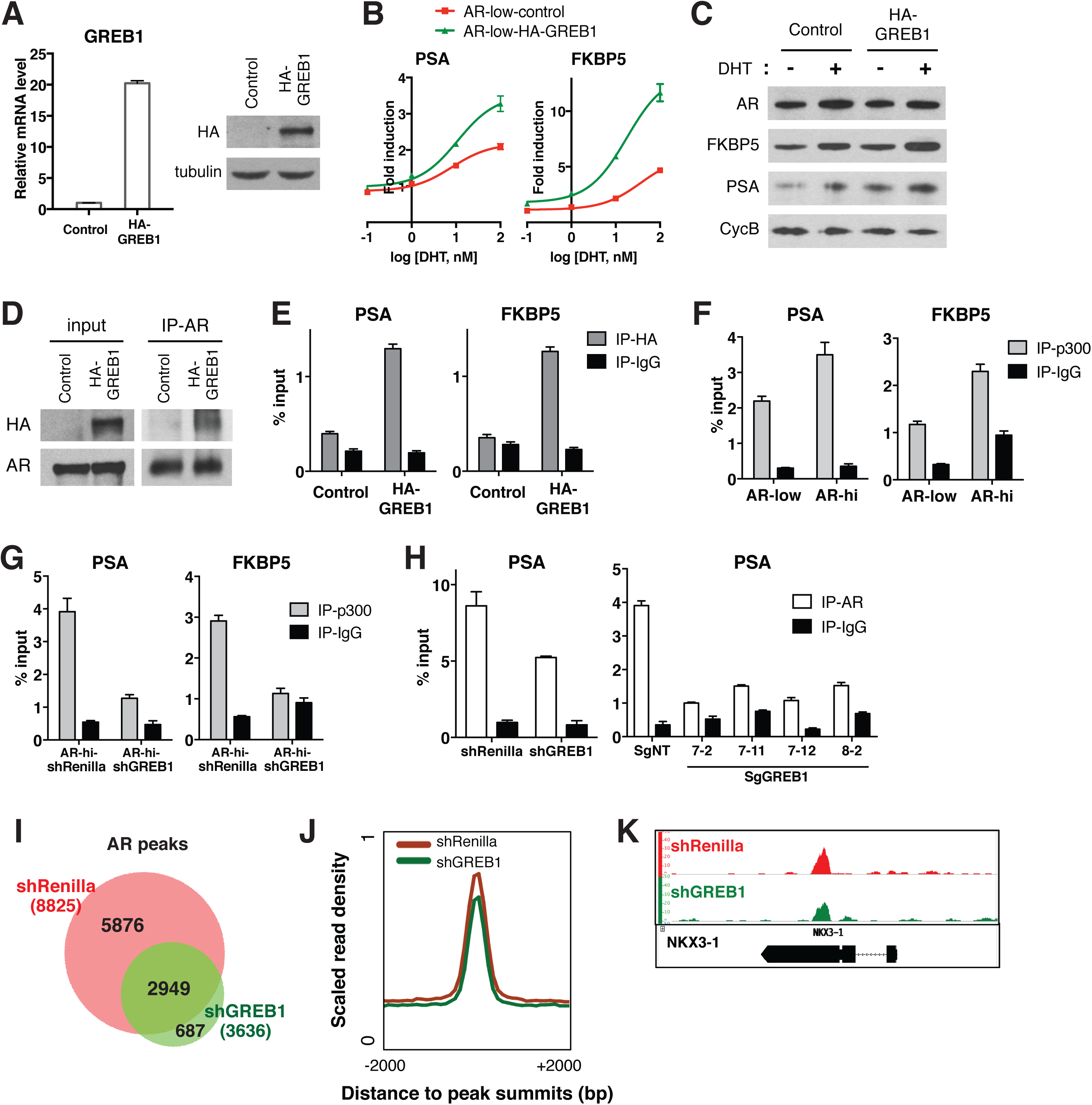
GREB1 amplifies AR transcriptional activity by enhancing AR binding to chromatin. (**A**) GREB1 overexpression in AR-low cells with stable integration of GREB1 vector containing HA-tag. (**B**) AR-low cells with GREB1 overexpression show higher induction of AR target genes in response to DHT treatment. The q-PCR data presented as mean fold change ± SD relative to DMSO control. (**C**) GREB1 overexpression in AR-low cells increases protein levels of AR target genes without affecting AR level. (**D**) Co-immunoprecipitation study using nuclear extract shows interaction between AR and GREB1 (HA). (**E**) ChIP against HA-tag shows GREB1 binding on PSA and FKBP5 enhancer regions. (**F-G**) AR-hi cells have increased p300 binding on PSA and FKBP5 enhancer regions in a GREB1 dependent manner. (**H**) GREB1 knockdown or CRISPR decreases AR binding to PSA enhancer in AR-hi cells. The ChIP q-PCR data presented as mean percentage input ± SD. (**I**) Overlap of AR ChIP-sequencing peaks shows that AR peaks are disrupted by GREB1 knocdown in AR-hi cells. (**J**) ChIP-sequencing summary plot shows that AR enrichment across the AR binding sites is reduced by GREB1 knockdown. (**K**) Example AR peaks on NKX3–1 gene.

In breast cancer, GREB1 functions as a coactivator through binding to ER and recruitment of the p300/CBP complex to ER target genes (Mohammed et al., 2013). We find that GREB1 functions similarly in prostate cells, as shown by co-immunoprecipitation documenting AR-GREB1 interaction (Figure 3D) and ChIP experiments showing recruitment of GREB1 to PSA and FKBP5 enhancer regions (Figure 3E). Furthermore, AR-hi cells showed a GREB1-dependent increase in p300 binding (Figure 3F,G) and GREB1 overexpression increased p300 recruitment to AR target genes in AR-low cells (Figure 3-figure supplement 2A).

In addition to this canonical coactivator function of promoting assembly of an active transcription complex, we found that GREB1 also impacts AR DNA binding. For example, knockdown or CRISPR deletion of GREB1 in AR-hi cells significantly reduced binding of AR to the PSA enhancer and, conversely, GREB1 overexpression promoted AR recruitment in AR-low cells (Figure 3H, Figure 3-figure supplement 2B). AR ChIP-sequencing revealed that this effect is genome-wide, with a significant reduction in the mean height of AR peaks in GREB1-depleted cells (Figure 3I-K). Importantly, the location of AR peaks (enhancer, promoter) was identical in intact versus GREB1 knockdown cells and there were no differences in consensus binding sites (Figure 3-figure supplement 2C,D). Therefore, GREB1 enhances AR DNA efficiency but not alter DNA binding site specificity. As seen previously in our analysis of AR-hi cells, total and nuclear AR levels were not changed by GREB1 knockdown or overexpression (Figure 3C, Figure 3-figure supplement 2E,F).

Of note, earlier studies of GREB1 in breast cancer did not report any effect on ER DNA binding (Mohammed et al., 2013), which we confirmed by GREB1 knockdown in MCF7 breast cancer cells (Figure 3-figure supplement 3A,B). Thus, GREB1 functions as a coactivator of both ER and AR but through somewhat different mechanisms. To address the possibility that other hormone receptor coactivators might also function differently in prostate cells, we asked if SRC-1 and SRC-2, previously shown to recruit the p300/CBP complex to AR (Leo & Chen, 2000), also influence AR DNA binding. To do so, we knocked down both genes in AR-hi cells based on prior work showing redundancy between SRC-1 and SRC-2 (Leo & Chen, 2000; Q. Wang, Carroll, & Brown, 2005). AR reporter activity and target gene expression was inhibited in SRC1/2-depleted cells, as expected, but AR occupancy of AR binding sites was unchanged (Figure 3-figure supplement 3C-E). Thus, in addition to a role in p300/CBP recruitment, GREB1 has unique effects on AR DNA binding that distinguish it from other coactivators.

### GREB1 is required for enzalutamide resistance of high AR output cells

Having demonstrated that GREB1 is overexpressed in AR-hi cells and functions as an AR coactivator, we asked if GREB1 is required for maintenance of the AR-hi state. First we evaluated the consequences of GREB1 knockdown on transcription. Consistent with experiments in AR-low cells showing that GREB1 overexpression enhanced AR transcriptional activity (Figure 3B,C, Figure 3-figure supplement 1B), GREB1 knockdown inhibited baseline and DHT-induced AR target gene expression in AR-hi cells (Figure 4A-C, Figure 4-figure supplement 1A,B). RNA-sequencing confirmed enrichment of androgen down-regulated gene sets in GREB1-depleted cells (Figure 4D) as well as downregulation of the 20 AR target genes used to calculate the AR activity score in TCGA tumors (Figure 4-figure supplement 1C). GREB1 knockdown cells also showed enrichment of the same prostate basal gene set that was enriched in AR-low cells (Figure 4D, refer also to Figure 2A). Additional analysis of RNA-seq data suggests that GREB1 is a major molecular determinant of the AR-hi state: specifically, (i) GREB1 knockdown impaired the induction of >70% of all DHT-induced genes (Figure 4E, Figure 4-source data 1,2) and (ii) the top 100 gene sets enriched in GREB1-depleted AR-hi cells and AR-low cells show significant overlap (Figure 4F, Figure 4-source data 3).

**Figure 4.**
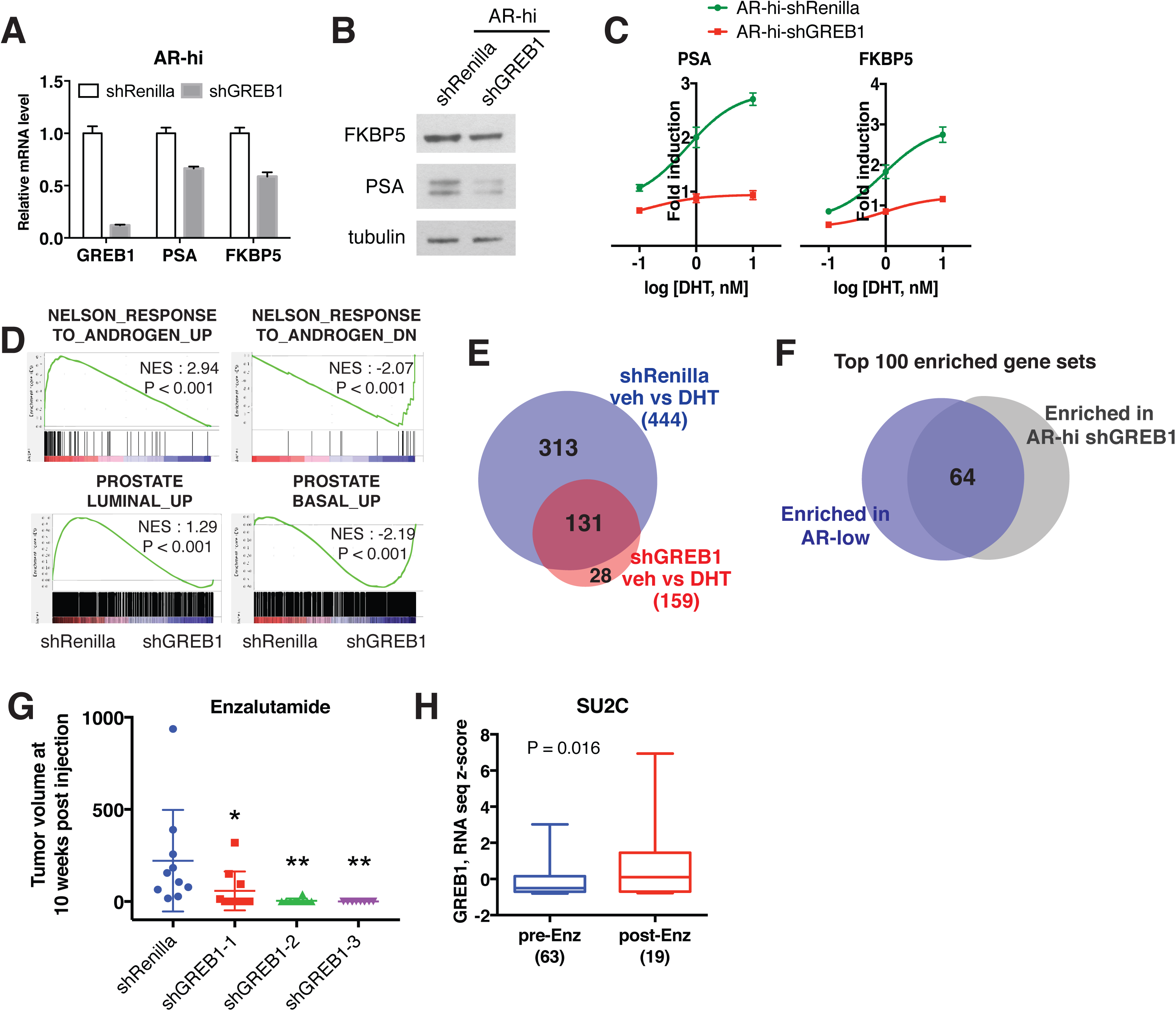
GREB1 is the major molecular determinant of AR-hi cells. **(A-B)** Knockdown of GREB1 inhibited AR target gene expression in AR-hi cells. The q-PCR data (A) presented as mean fold change ± SD relative to shRenilla control. (**C**) Knockdown of GREB1 suppressed the enhanced AR transcriptional activity in AR-hi cells. The q-PCR data presented as mean fold change ± SD relative to DMSO control. (**D**) Gene set enrichment analysis (GSEA) shows that the gene sets up- and down-regulated by androgen are enriched in AR-hi control and GREB1 knockdown cells, and genes upregulated in human prostate luminal and basal cells are enriched in AR-hi control and GREB1 depleted cells, respectively. (**E**) The venn diagram showing that 70.5% of DHT-induced genes in control AR-hi cells was inhibited by GREB1 knockdown. (**F**) The venn diagram showing that 64% of top 100 gene sets enriched in AR-low overlaps with top 100 gene sets enriched in GREB1 depleted AR-hi cells. (**G**) Knockdown of GREB1 inhibited development of enzlutamide-resistant LNCaP/AR xenografts derived from AR-hi cells. The sorted AR-hi cells were infected with control or 3 different shRNAs targeting GREB1 and injected into castrated mice. Mice were treated with enzalutamide immediately after injection. Data presented as mean ± SEM (N=10). *P<0.05, **P<0.01 (One-way ANOVA). (**H**) The SU2C cases received enzalutamide (Enz) have increased level of GREB1 (unpaired t-test).

Earlier we showed that AR-hi cells rapidly acquire resistance to enzalutamide (refer to Figure 1I). To determine the role of GREB1 in this drug resistant phenotype, we performed knockdown experiments using the LNCaP/AR xenograft. After confirming that AR activity was inhibited in AR-hi cells (Figure 4-figure supplement 1D,E), we injected LNCaP/AR-hi xenografts with GREB1 shRNAs into castrated mice treated with enzalutamide and found a significant delay in the development of enzalutamide resistance after 10 weeks (Figure 4G). Clinical data from CRPC patients also supports for a role of GREB1 in enzalutamide resistance, based on increased GREB1 expression in those who progressed on enzalutamide treatment (Figure 4H).

## Discussion

There is abundant evidence from tumor sequencing studies that genomic alterations in AR (amplification and/or mutation) are present in over 50% of CRPC patients (Cancer Genome Atlas Research, 2015; Robinson et al., 2015) and that AR amplification is associated with a less favorable clinical response to abiraterone or enzalutamide treatment (Annala et al., 2018; Podolak et al., 2017). Therefore, high levels of AR transcriptional output can promote castration-resistant disease progression. Here we show that prostate cancers can amplify AR output through increased expression of the dual AR/ER coactivator GREB1, in the absence of genomic AR alterations. As with genomic AR amplification, increased AR output driven by high GREB1 expression is also associated with enzalutamide resistance.

In addition to demonstrating the importance of transcriptional heterogeneity in drug resistance, we also show that GREB1 amplifies AR activity by a novel two-part mechanism. Similar to canonical coactivators such as SRC1/2, GREB1 binds AR and promotes the assembly of an active transcription complex by recruitment of histone acetyl transferases such as p300/CBP (Lee & Lee Kraus, 2001). However, GREB1 has the additional property of improving the efficiency of AR binding to DNA, which further enhances AR transcriptional output. Although conceptually distinct from canonical coactivators, this dual mechanism of AR activation is may not be unique to GREB1. For example, TRIM24 has been shown to function as an oncogenic AR cofactor and, similar to GREB1, knockdown of TRIM24 impairs recruitment of AR to target genes (Groner et al., 2016). Curiously, the effect of GREB1 on AR DNA binding is not seen with ER, suggesting different conformational consequences of GREB1 binding on AR and ER respectively then influence DNA binding.

One curious observation is the fact that prostate cancers can maintain transcriptional heterogeneity as a stable phenotype, despite the fact that GREB1 expression drives a feed forward loop which, in principle, should result in an increased fraction of high AR output cells over time. One potential explanation for the ability of these populations to maintain stable proportions of high versus low AR output cells at steady state is the fact that androgen has growth inhibitory effects at higher concentrations (Culig et al., 1999). Because GREB1 amplifies the magnitude of AR output in response to normal (growth stimulatory) androgen concentrations, the biologic consequence of high GREB1 levels could be the same growth suppression seen with high androgen concentrations. This model predicts that high AR output cells would gain a fitness advantage under conditions of androgen deprivation or pharmacologic AR inhibition, as demonstrated by the enzalutamide resistance observed in xenograft models.

Further work is required to understand the clinical implications of our work but two findings deserve comment. First, we show that elevated levels of GREB1 in CRPC tumors correlate with a poor clinical response to enzalutamide, analogous to the prognostic impact of AR gene amplification. Second, GREB1 knockdown experiments provide genetic evidence that GREB1 is required for in vivo enzalutamide resistance in xenograft models. Although pharmacologic strategies to inhibit GREB1 function are not currently available, a small molecule inhibitor that blocks protein-protein interactions between the AR N-terminal domain and CBP/p300 is currently in clinical development (Andersen et al., 2010) (NCT02606123). This work provides precedent that similar strategies to disrupt GREB1/AR interaction may be possible.

## Materials and Methods

### Cell lines

LNCaP and MCF7 cell lines were obtained from American Type Culture Collection (ATCC, Manassas, VA) and maintained in RPMI (LNCaP) or DMEM (MCF7) + 10% FBS (Omega Scientific, Tarzana, CA). LNCaP/AR cell line was generated and maintained as previously described (C. D. Chen et al., 2004). CWR22Pc was a gift from Marja T. Nevalainen (Thomas Jefferson University, Philadelphia, PA) and CWR22Pc-EP was generated and maintained as previously described (Mu et al., 2017).

### Flow cytometry analysis and FACS-sorting

Rapidly cycling eGFP AR reporter cells were collected using Accumax dissociation solution (Innovative Cell Technologies, San Diego, CA), and dead cells were counterstained with DAPI (Invitrogen, Grand Island, NY). For FACS-sorting of AR-low and AR-hi cells, 5% of the entire population with lowest and highest eGFP expression was sorted out using BD FACSAria cell sorter. For flow cytometric analysis of reporter activity, eGFP expression was measured using the BD-LDRII flow cytometer and analysis was done using FlowJo software.

### Plasmid construction and cell transduction

The lentiviral eGFP AR reporter (ARR3tk-eGFP/SV40-mCherry) was generated by switching 7xTcf promoter of 7xTcf-eGFP/SV40-mCherry (Addgene, Cambridge, MA, 24304) with probasin promoter containing 3xARE (ARR_3_tk) (Snoek et al., 1998). For shRNA knockdown experiments, SCEP vector was generated by substituting GFP cassette of SGEP (pRRL-GFP-miRE-PGK-PuroR, gift from Johannes Zuber) (Fellmann et al., 2013) with mCherry cassette. The following guide sequences were used for knockdown:

shAR.177: TAGTGCAATCATTTCTGCTGGC

shGREB1–1: TTGTCAGGAACAGACACTGGTT

shGREB1–2: TTTCAGATTTATATGATTGGAG

shGREB1–3: TTGACAAGATACCTAAAGCCGA

shKLF8.3467: TTGAGTTCTAAAGTTTTCCTGA

shKLF8.2180: TATTTGTCCAAATTTAACCTAA

shKLF8.2684: TTATAAAACAATCTGATTGGGC

shGHRHR.544: TAAAAGTGGTGAACAGCTGGGT

shGHRHR.1571: TTTATTGGCTCCTCTGAGCCTT

shGHRHR.1583: TTCATTTACAGGTTTATTGGCT

shSRC1–1: TTCTTCTTGGAACTTGTCGTTT

shSRC2–1: TTGCTGAACTTGCTGTTGCTGA

shSRC2–2: TTAACTTTGCTCTTCTCCTTGC

shRenilla was previously described as Ren.713 targeting Renilla luciferase (Fellmann et al., 2013). Pools of 3 shRNAs were used to knockdown GREB1, KLF8 and GHRHR in a small-scale shRNA screen, and shGREB1–1 was used for further studies. For CRISPR/Cas9 experiments, lentiCRISPRv2 vector gifted by F. Zhang (Addgene, 52961) was used with the following guide sequences designed using http://crispr.mit.edu/website:

SgGREB1–7: AGGCATGTCCTGCGTGCCGC

SgGREB1–8: TCACGGGCATACGAGCAGTA

sgNT was previously described (T. Wang, Wei, Sabatini, & Lander, 2014). pCMV6-GREB1 plasmid was a gift from J. Carroll (Cancer Research UK Cambridge Institute, Cambridge, UK). The lentiviral GREB1 cDNA plasmid was constructed by cloning GREB1 cDNA from pCMV6-GREB1 into Tet-inducible pLV-based lentiviral expression vector with HA-tag.

Lentiviral transduction of cells was performed as described previously (Mu et al., 2017). To make AR reporter cell line, cells were infected with ARR3tk-eGFP/SV40-mCherry at low multiplicity of infection (MOI) to enable each cell has one copy of reporter construct, and the transduced cells were sorted by mCherry flow cytometry. To inactivate GREB1 gene, we single-cell cloned the cells infected with lentiCRISPRv2 vector containing SgGREB1–7 or SgGREB1–8, and isolated a clone that had genomic alteration at target sequence. Three clones were generated by using SgGREB1–7 (SgGREB1–7–2, 7–11 and 7–12) and one clone was generated by using SgGREB1–8 (SgGREB1–8–2).

### shRNA screen

FACS-based small-scale shRNA screen with 33 selected genes was performed as follows: FACS-sorted AR-hi cells were plated in 12 well plate (1.5 × 10^5^ cells per well, Corning, 353043) and each well was infected with pool of 3 SEPC shRNAs against each gene on the following day. Cells with stable integration of hairpins were selected with 2 μg/ml puromycin. 9 days after infection, half of the cells in each well was used to analyze eGFP AR reporter activity using flow cytometry, and the other half was subjected to qRT-PCR to determine knockdown level of the gene. We performed the screen in duplicate and each replicate included wells infected with shRenilla or shAR as controls. The median fluorescence intensity (MFI) of eGFP was measured using FlowJo software. The shRNAs decreased eGFP MFI more than 1.5 fold compared to shRenilla (normalized value lower than 0.667) in both duplicate were considered as hits. The list of 33 genes used in the screen and the summary of median eGFP intensity can be found at Figure2-source data 2.

### Xenograft assay

To compare time to acquire enzalutamide resistance *in vivo*, FACS-sorted bulk, AR- low and AR-hi populations derived from LNCaP/AR were cultured for 6 days after sorting to obtain enough number of cells for xenograft assay. 2 × 10^6^ cells were injected subcutaneously into the flank of castrated CB17 SCID mice in a 50:50 mix of matrigel (BD Biosciences, San Jose, CA) and regular culture medium (5 mice, 10 tumors per group), and enzalutamide treatment was initiated on the day of injection. To test the effect of GREB1 knockdown on development of enzalutamide resistance, FACS-sorted AR-hi population derived from LNCaP/AR was infected with control or 3 different shGREB1 constructs 2 days after sorting. Cells with stable integration of hairpin were selected with 2 μg/ml puromycin. 5 days after infection, 2 × 10^6^ cells were injected subcutaneously into the flank of castrated CB17 SCID mice (5 mice, 10 tumors per group), and enzalutamide treatment was initiated on the day of injection. The same cell populations used for injection were also used to test eGFP AR reporter activity using flow cytometry, and qRT-PCR to test knockdown level of GREB1. Measurements were obtained weekly using Peira TM900 system (Peira bvba, Belgium). All animal experiments were performed in compliance with the guidelines of the Research Animal Resource Center of the Memorial Sloan Kettering Cancer Center.

### Immunoblot, immunoprecipitation and immunostaining

Protein was extracted from cells using Triton lysis buffer and quantified by BCA Protein Assay (ThermoFisher Scientific, Waltham, MA, 23225). Nuclear/cytoplasmic fractionation was achieved with Subcellular Protein Fractionation Kit (ThermoFisher Scientific, 78840). Protein lysates were subjected to SDS-PAGE and immunoblotted with the following antibodies against: AR (Abcam, Cambridge, United Kingdom, ab108341), PSA (Cell Signaling Technology, Danvers, MA, 5365), FKBP5 (Cell Signaling, 8245) TRPM8 (Epitomics, Burlingame, CA, 3466–1), tubulin (Santa Cruz Biotechnology, Dallas, TX, sc-9104), Cyclophilin B (Abcam, ab178397), BRD4 (Cell Signaling, 13440), TOP2B (Abcam, ab58442), HA (Cell Signaling, 3724).

For AR immunoprecipitation, at least 1.5 mg of total protein was incubated with AR antibody (Abcam, ab108341) overnight at 4 °C followed by the addition of Protein A/G agarose beads (Santa Cruz, sc-2003) for 2 h. Immune complexes were extensively washed with Triton buffer and solubilized using Laemmli sample buffer (BioRad, Hercules, CA).

For immunofluorescence staining, cells were fixed with 4% formaldehyde, permeabilized with 0.2% Triton-X, blocked with 5% normal goat and 5% normal horse serum, stained with anti-AR (Santa Cruz, sc-816) primary and Alexa Fluor 647 (Invitrogen) secondary antibodies, and mounted with DAPI mounting solution (Vector Lab, Burlingame, CA). For Immunohistochemistry, tumor sections were stained with anti-AR (Agilent, Santa Clara, CA, 441) and PSA (Biogenex, Fremont, CA) antibodies using Leica Bond RX (Leica Biosystems, Wetzlar, Germany).

### Transcription analysis

Total RNA was isolated using the QiaShredder kit (Qiagen, Germantown, MD) for cell lysis and the RNeasy kit (Qiagen) for RNA purification. For quantitative PCR with reverse transcription (RT–qPCR), we used the High Capacity cDNA Reverse Transcription Kit (Applied Biosystems, Grand Island, NY) to synthesize cDNA according to the manufacturer’s protocol. Real-time PCR was performed using gene-specific primers and 2X SYBR green quantfast PCR Mix (Qiagen, 1044154). Data were analyzed by the DDCT method using GAPDH as a control gene and normalized to control samples, which were arbitrarily set to 1. To test DHT-induced AR target gene upregulation, cells were hormone-deprived in 10% charcoal-stripped dextran-treated fetal bovine serum (Omega Scientific) media for 2 days and then treated with indicated concentration of DHT for 24 h. Triplicate measurements were made on at least three biological replicates. The primer sequences used for q-PCR are listed at Supplementary file 1.

For RNA-seq, library preparation, sequencing and expression analysis were performed by the New York Genome Center. Libraries were prepared using TruSeq Stranded mRNA Library Preparation Kit in accordance with the manufacturer’s instructions and sequenced on an Illumina HiSeq2500 sequencer (rapid run v2 chemistry) with 50 base pair (bp) reads. Partek^®^ Genomics Suite^®^ software (Partek Inc, St. Louis, MO) was used to analyze differentially expressed genes between AR-low vs. AR-hi (Fold change ≥ 1.5, p < 0.05). To analyze RNA-seq data from AR-hi cells with shRenilla vs. shGREB1, reads were aligned to the NCBI GRCh37 human reference using STAR aligner (Dobin et al., 2013). Quantification of genes annotated in Gencode vM2 were performed using featureCounts and quantification of transcripts using Kalisto (Bray, Pimentel, Melsted, & Pachter, 2016). QC were collected with Picard and RSeQC (L. Wang, Wang, & Li, 2012) (http://broadinstitute.github.io/picard/). Normalization of feature counts was done using the DESeq2 package (http://www-huber.embl.de/users/anders/DESeq/). Differentially expressed genes were defined as a 1.5 fold difference, p < 0.05 of DESeq-normalized expression. For GSEA, statistical analysis was performed with publicly available software from the Broad Institute (http://www.broadinstitute.org/gsea/index.jsp). The basal and luminal gene signatures used for GSEA (Supplementary file 2) were generated by conducting RNA-sequencing with normal human basal vs. luminal prostate cells isolated as previously described (Karthaus et al., 2014). Full description of this study will be reported separately.

### ChIP

ChIP experiments were performed as previously described (Arora et al., 2013), using SDS-based buffers. Antibodies were used at a concentration of 5 ug per 1 mL of IP buffer, which encompassed approximately 8 million cells per IP. Antibodies used were: AR (Santa Cruz, sc-816), p300 (Santa Cruz, sc-585), HA (Abcam, ab9110), ER (Santa Cruz, sc-8002). The primer sequences used for ChIP-qPCR are listed at Supplementary Table S8.

For ChIP–seq, library preparation and RNA-seq were performed by the NYU Genome Technology Center. Libraries were made using the KAPA Biosystems Hyper Library Prep Kit (Kapa Biosystems, Woburn, MA, KK8504), using 10 ng of DNA as input and 10 PCR cycles for library amplification. The libraries were sequenced on a HiSeq 2500, as rapid run v2 chemistry, paired-end mode of 51 bp read length.

The ChIP-seq reads were aligned to the human genome (hg19, build 37) using the program BWA (VN: 0.7.12; default parameters) within the PEMapper. Duplicated reads were marked by the software Picard (VN: 1.124; http://broadinstitute.github.io/picard/index.html) and removed. The software MACS2 (Feng, Liu, Qin, Zhang, & Liu, 2012) (-q 0.1) was used for peak identification with data from ChIP input DNAs as controls. Peaks of sizes >100 bp and with at least one base pair covered by >18 reads were selected as the final high confident peaks. Peaks from shGREB1/control conditions were all merged to obtain non-overlapping genomic regions, which were then used to determine conditional specific AR binding. Overlapped peaks were defined as those sharing at least one base pair. To generate graphs depicting AR ChIP–seq read density in ±2 kilobase regions of the AR peak summits, the same number of ChIP–seq reads from different conditions were loaded into the software ChAsE (Younesy et al., 2016), and the resulting read density matrices were sorted by the read densities in the shRenilla control, before colouring. The read density was also used to select peaks with significant signal difference between shGREB1 and controls. The criteria for assigning peaks to genes have been described previously (Rockowitz & Zheng, 2015). The MEME-ChIP software (Machanick & Bailey, 2011) was applied to 300-bp sequences around the peak summits for motif discovery, and the comparison of sequence motifs was also analyzed with HOMER (http://homer.ucsd.edu/homer/).

### Analysis of human prostate cancer datasets

All analysis of human prostate cancer data was conducted using previously published datasets of The Cancer Genome Atlas (TCGA) (Cancer Genome Atlas Research, 2015) and PCF/SU2C (Robinson et al., 2015), which can be explored in the cBioPortal for Cancer Genomics (http://www.cbioportal.org).

### Statistics

For comparison of pooled data between two different groups, unpaired t tests were used to determine significance. For comparison of data among three groups, one-way ANOVA was used to determine significance. In vitro assays represent three independent experiments from biological replicates, unless otherwise indicated. In all figures, \**P*<0.05, \*\**P*<0.01 and \*\*\**P*<0.001. For GSEA, statistical analysis was performed with publicly available software from the Broad Institute (http://www.broadinstitute.org/gsea/index.jsp). The sample size estimate was based on our experience with previous experiments (Balbas et al., 2013; Bose et al., 2017; Y. Chen et al., 2013). No formal randomization process was used to assign mice to a given xenograft assay, and experimenters were not blinded.

## Acknowledgments

We thank the flow cytometry core facility at MSKCC for technical support, NYU Genome Technology Center for conducting ChIP-sequencing, New York Genome Center for conducting the RNA-sequencing, MSKCC Pathology Core for assistance with IHC staining of patient samples, Wouter Karthaus for help with cloning and providing the basal and luminal gene signature, Wassim Abida for help with analyzing patient data, Kayla Lawrence and Tejasveeta Nadkarni for help with cloning and Jason Carroll (Cancer Research UK Cambridge Institute) for generously providing pCMV6-GREB1 plasmid, and the members of the Sawyers laboratory for helpful discussions.

## Disclosure Statement

Charles L Sawyers: Senior Editor eLife; Board of Directors of Novartis; co-founder of ORIC Pharm; co-inventor of enzalutamide and apalutamide; Science advisor to Agios, Beigene, Blueprint, Column Group, Foghorn, Housey Pharma, Nextech, KSQ, Petra and PMV; co-founder of Seragon, purchased by Genentech/Roche in 2014. John Wongvipat is a co-inventor of enzalutamide.

---

**Figure 1-figure supplement 1.**
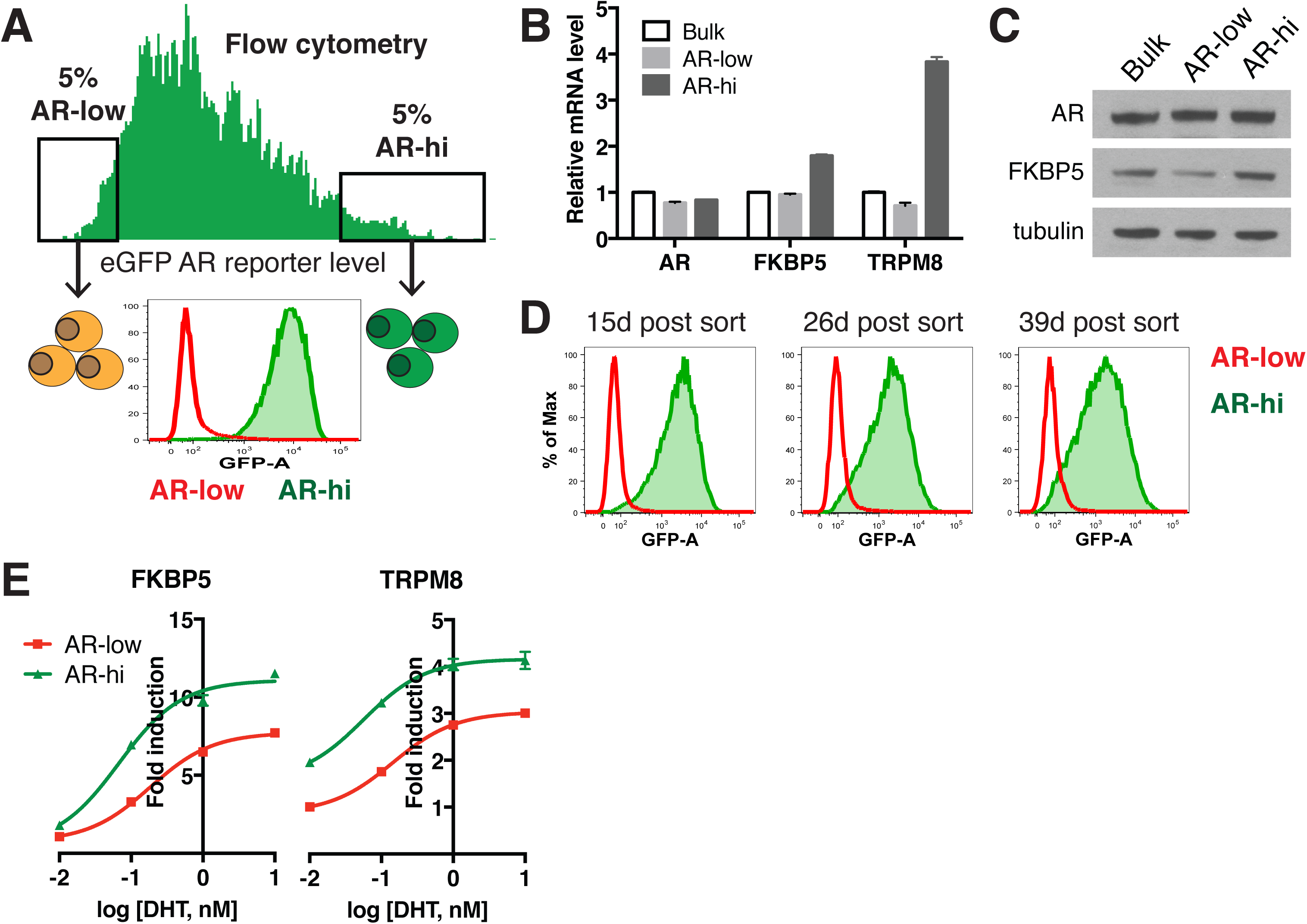
Characterization of CWR22Pc-EP prostate cancer cells with low vs. high AR output. (**A**) CWR22Pc-EP cells with low (AR-low) and high (AR-hi) AR activities were sorted out using flow cytometry based on eGFP AR-reporter expression. (**B-C**) CWR22Pc-EP AR-hi cells have higher AR output while having same level of AR. The q-PCR data (B) presented as mean fold change ± SD relative to bulk population. (**D**) CWR22Pc-EP AR-low and AR-hi cells maintain their AR activities over time. (**E**) CWR22Pc-EP AR-hi cells have enhanced DHT-induced AR transcriptional activity compared to AR-low cells. The q-PCR data presented as mean fold change ± SD relative to DMSO control.

**Figure 1-figure supplement 2.**
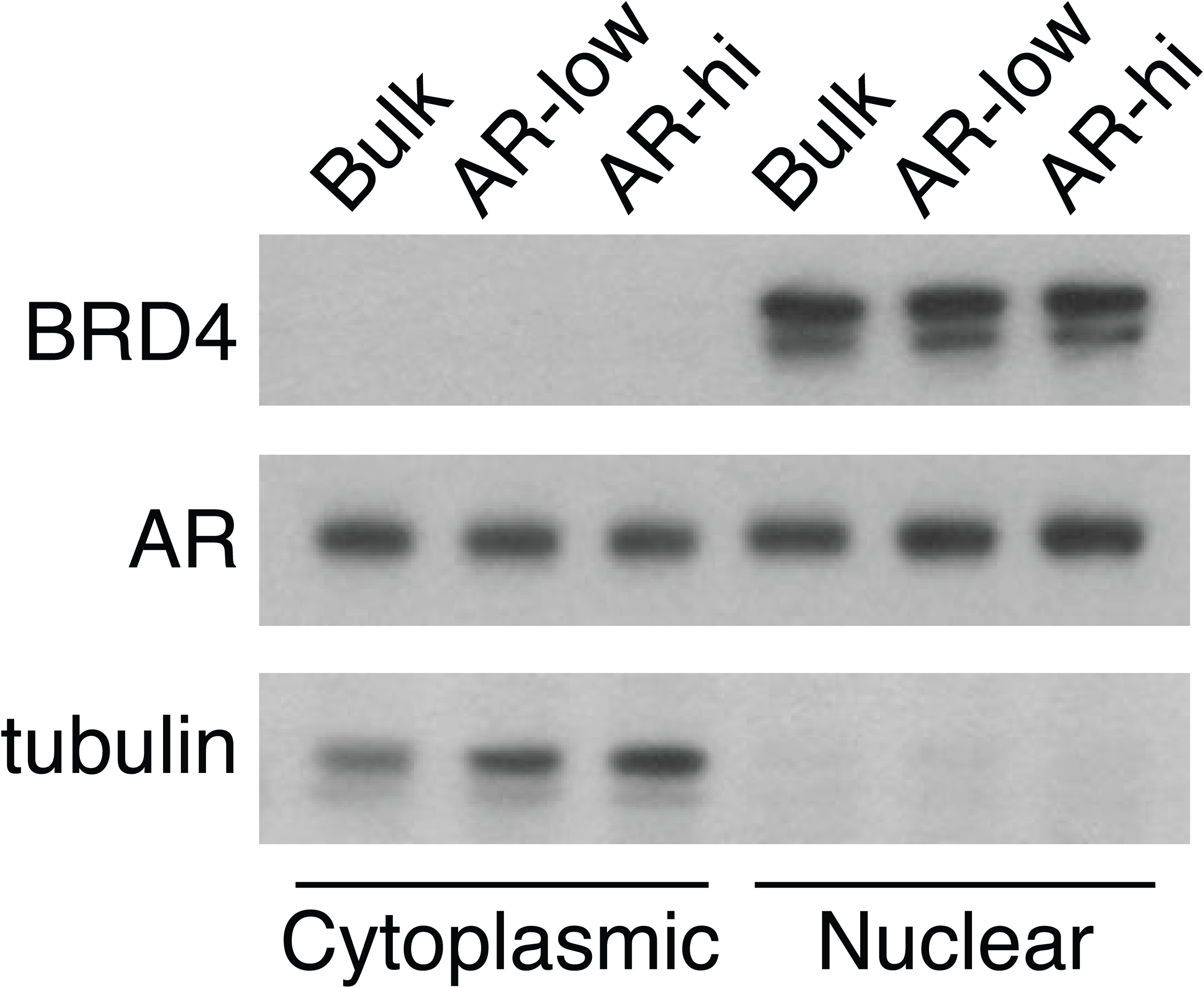
LNCaP AR-low and AR-hi cells have comparable nuclear AR level. The BRD4 and tubulin were used as nuclear and cytoplasmic loading controls, respectively.

**Figure 1-figure supplement 3.**
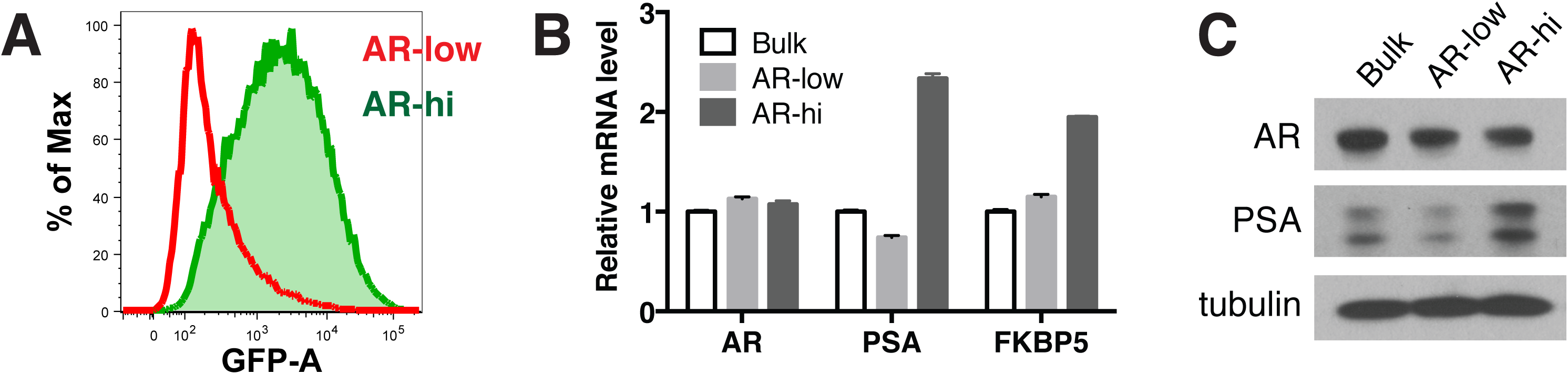
Characterization of LNCaP/AR prostate cancer cells with low vs. high AR output. (**A-C**) LNCaP/AR cells with low (AR-low) and high (AR-hi) AR activities were sorted out using flow cytometry based on eGFP AR-reporter expression. The AR reporter activity (A) and AR target gene expression (B-C) were analyzed 7 days post sorting.

**Figure 1-figure supplement 4.**
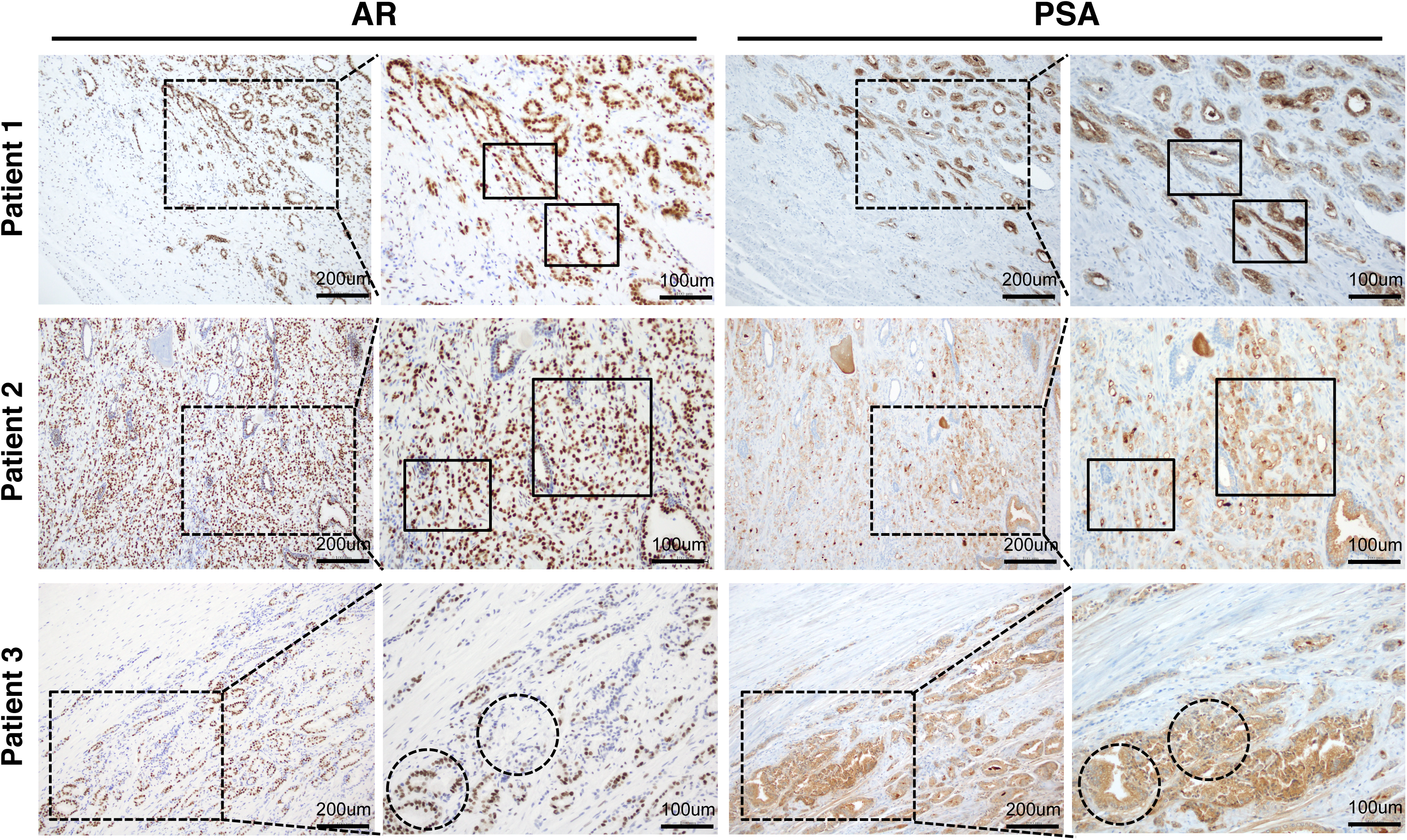
AR and PSA staining in untreated localized prostate cancer shows heterogeneous PSA staining that is not strictly correlated with AR level.

**Figure 2-figure supplement 1.**
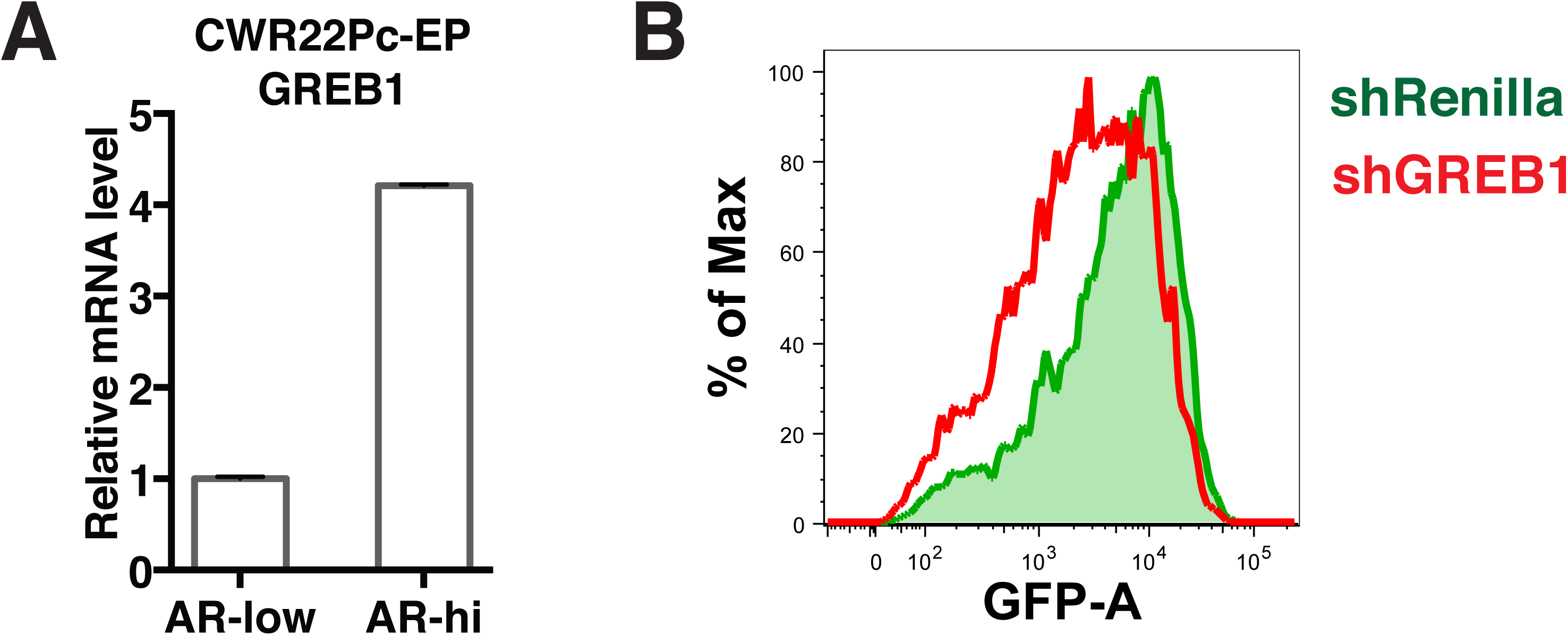
Inhibition of GREB1 suppresses AR transcriptional activity in CWR22Pc-EP cells. (**A**) GREB1 is upregulated in CWR22Pc-EP AR-hi cells. (**B**) Knockdown of GREB1 in CWR22Pc-EP AR-hi cells inhibited AR reporter activity.

**Figure 3-figure supplement 1.**
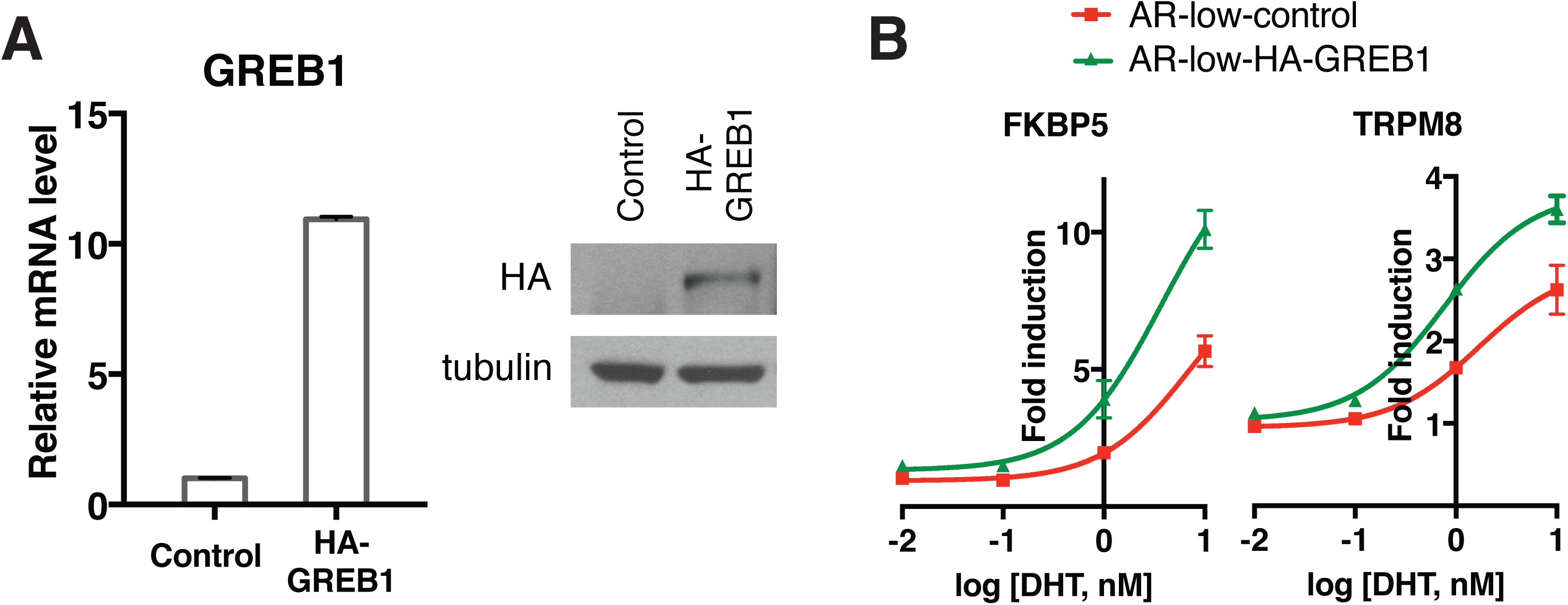
GREB1 amplifies AR transcriptional activity in CWR22Pc-EP cells. (**A**) GREB1 overexpression in AR-low cells with stable integration of GREB1 vector containing HA-tag. (**B**) AR-low cells with GREB1 overexpression show higher induction of AR target genes in response to DHT treatment. The q-PCR data presented as mean fold change ± SD relative to DMSO control.

**Figure 3-figure supplement 2.**
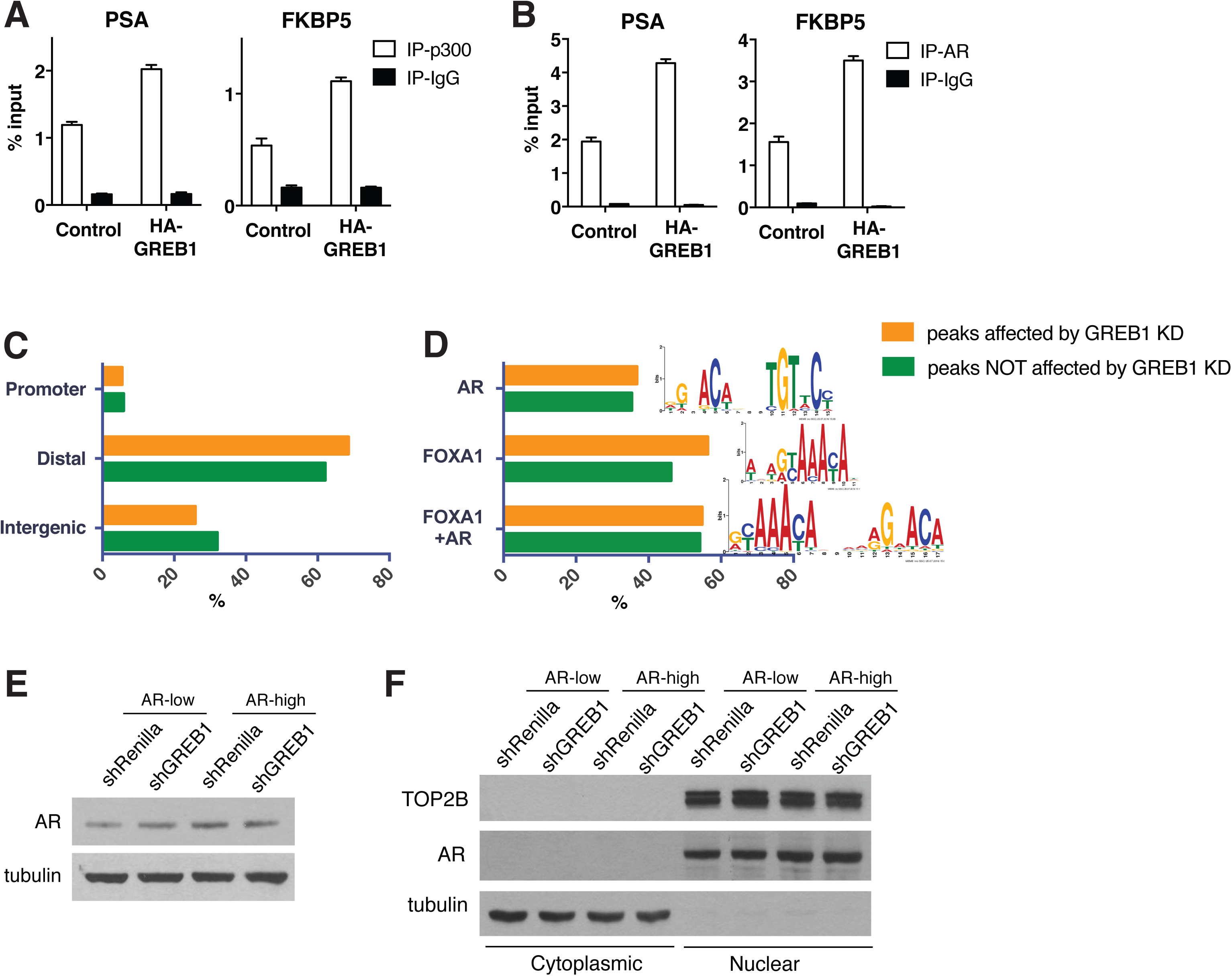
GREB1 enhances AR binding to chromatin. **(A-B)** GREB1 overexpression promotes p300 (A) and AR (B) binding to PSA and FKBP5 enhancer regions in LNCaP AR-low cells. The ChIP q-PCR data presented as mean percentage input ± SD. (**C-D**) The location (C) and motif (D) analysis of the AR ChIP-sequencing data shows no difference between peaks affected and not affected by GREB1 knockdown. (**E-F**) GREB1 knockdown has no effect on total (E) and nuclear (F) AR level in both LNCaP AR-low and AR-hi cells. The BRD4 and tubulin were used as nuclear and cytoplasmic loading controls, respectively.

**Figure 3-figure supplement 3.**
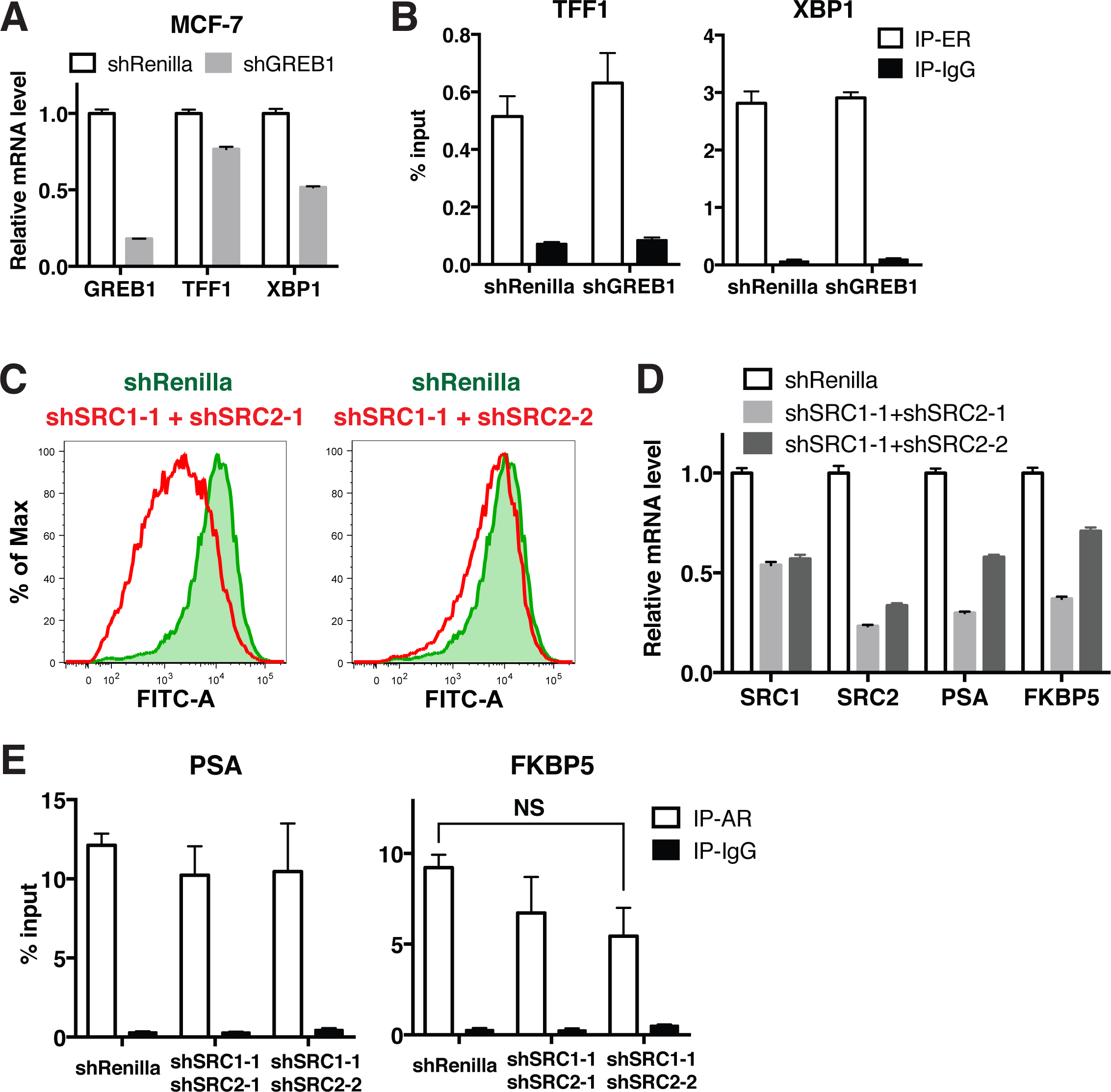
GREB1 has unique function compared to ER or SRC-1 and SRC-2. (**A-B**) Knockdown of GREB1 in MCF7 cells inhibits ER target gene expression (A), but has no affect on ER recruitment to binding sites (B). (**C-E**) Knockdown of both SRC1 and SRC2 in LNCaP AR-hi cells inhibits AR reporter activity (C) and AR target gene expression (D), but does not affect AR binding on PSA and FKBP5 enhancer regions (E).

**Figure 4-figure supplement 1.**
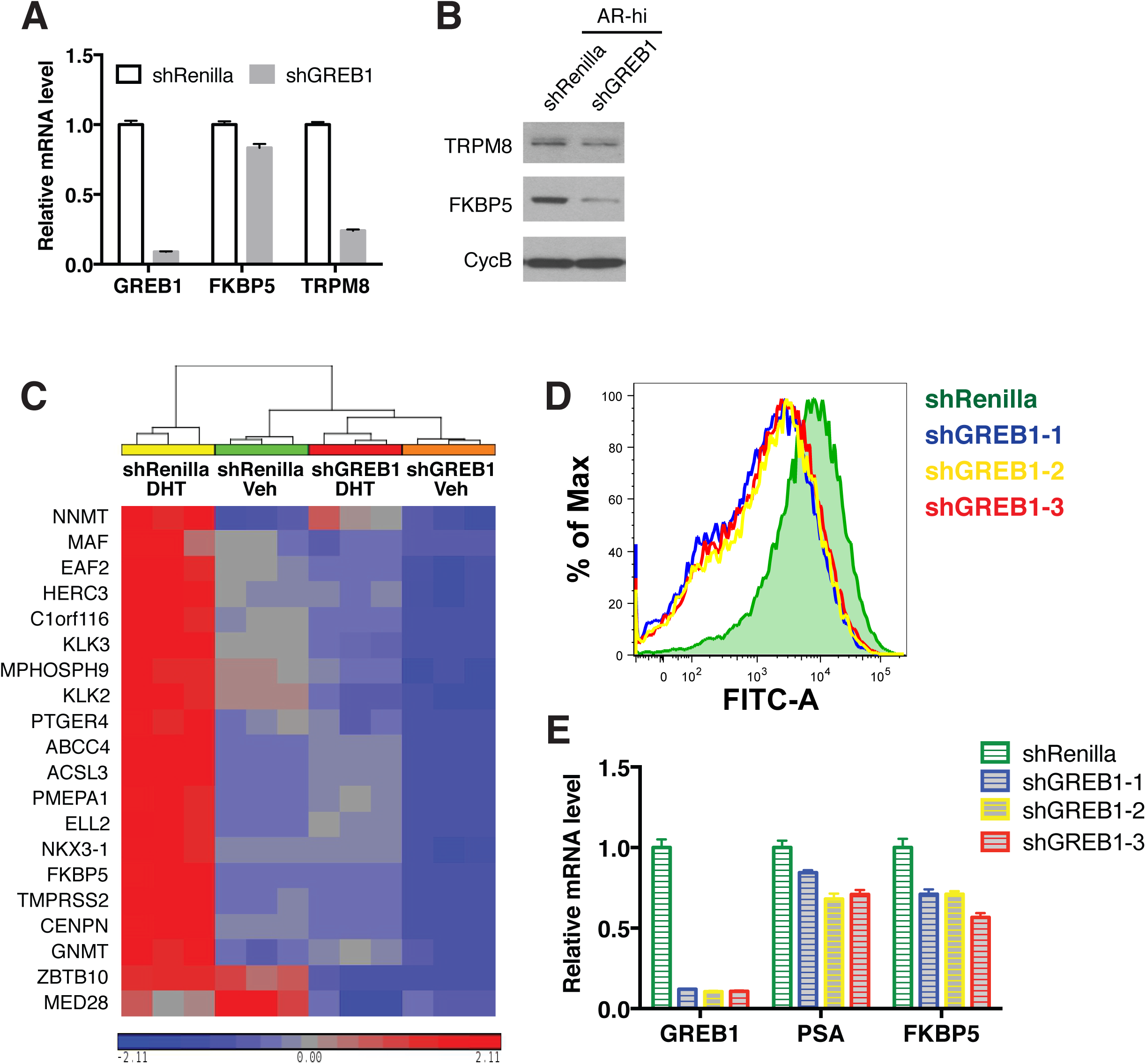
Knockdown of GREB1 inhibits AR signaling. (**A-B**) Knockdown of GREB1 inhibited AR target gene expression in CWR22Pc-EP AR-hi cells. (**C**) The heatmap generated from the RNA-sequencing data shows that the expression of 20 AR target genes used to calculate AR score is suppressed by GREB1 knockdown in LNCaP AR-hi cells. (**D-E**) Knockdown of GREB1 in AR-hi cells derived from LNCaP/AR inhibited AR reporter activity (D) and expression of AR target genes (E).

## Source data and Supplementary files

**Figure 1-source data 1**: GSEA Results (AR-low vs. AR-hi)

**Figure 2-source data 1**: Differentially expressed genes between AR-low vs. AR-hi

**Figure 2-source data 2**: Summary of Median eGFP Intensity of small-scale shRNA screen

**Figure 4-source data 1**: Upregulated genes in AR-hi shRenilla DHT vs. veh

**Figure 4-source data 2**: Upregulated genes in AR-hi shGREB1 DHT vs. veh

**Figure 4-source data 3**: GSEA Results (AR-hi shRenilla DHT vs. shGREB1 DHT)

**Supplementary file 1**: Primer list

**Supplementary file 2**: The basal and luminal gene signatures used for GSEA

